# Pan-EphB tyrosine kinase inhibitor mitigates diabetic cardiomyopathy

**DOI:** 10.1101/2025.11.17.688935

**Authors:** Heba A. Ewida, Md Tareq Aziz, Syed Tareq, Prasanth K. Chelikani, Mahmoud Salama Ahmed

**Affiliations:** Department of Pharmaceutical Sciences, Jerry H. Hodge School of Pharmacy, Texas Tech University Health Sciences Center, Amarillo, TX 79106, USA; School of Veterinary Medicine, Texas Tech University, Amarillo, TX, 79106, USA; Brain Drug Discovery Center, Texas Tech University Health Sciences Center, Amarillo, TX 79106, USA

## Abstract

Diabetic cardiomyopathy (DCM) is characterized by myocardial fibrosis, excessive collagen deposition, decline in ejection fraction and diastolic dysfunction in diabetic subjects. As current pharmaceuticals have varying efficacy and adverse effects, there is an unmet need for new therapeutics. We recently identified STA-013, a pan-EphB tyrosine kinase inhibitor sparing EphB3, that resulted in whole-body weight loss, restored glucose homeostasis, and mitigated insulin resistance in diet-induced obese mouse models. Here, we now show in a streptozotocin (STZ)-high-fat diet-induced mouse model that STA-013 decreased body weight and restored glucose homeostasis while mitigating insulin resistance. Importantly, STA-013 significantly improved cardiac function in part by increasing ejection fraction and fractional shortening, and reduced collagen deposition suggestive of decreased cardiac fibrosis. This was associated with a significant downregulation of key molecular intermediaries including phosphorylation EphB tyrosine kinase signaling, insulin receptor signal, and TGF-β levels in STA-013 treated STZ-HFD mice. Together, our preclinical findings support the development of pan-EphB tyrosine kinase inhibitors as novel therapeutics to mitigate DCM and associated complications.

## Introduction

**Diabetic cardiomyopathy (DCM)** is defined by the presence of uncontrolled Type-2 diabetes (T2D) that can lead to impaired cardiac insulin signaling associated with decreased mitochondrial functions, increased oxidative stress signaling, myocardial fibrosis, and excessive collagen deposition. The collagen will stiffen the heart tissue, leading to dysfunctional remodeling, heart failure with reduced ejection fraction, diastolic dysfunction, and cardiomyopathy^1–4^. The main underlying molecular mechanisms associated with the progression of DCM involve crosstalk among multiple molecular signature proteins including AMP-activated protein kinase, peroxisome proliferator-activated receptors, O-linked N-acetylglucosamine, nuclear factor erythroid 2–related factor 2 (Nrf2). protein kinase C, sodium-glucose co-transporter-II (SGLT-2), microRNA, and exosome pathways^5^. Current therapeutic strategies for patients living with DCM include lipid-lowering drugs, traditional anti-diabetics, exercise-promoting lifestyle changes, and conventional heart failure (HF) medications, which include β Blockers, angiotensin-converting enzyme inhibitors/angiotensin-receptor blockers (ACEI/ARBs), and SGLT-2 inhibitors^6^. Further, glucagon-like peptide 1 receptor agonists (semaglutide) reduced cardiovascular events and heart failure in high-risk diabetic patients^7,8^. However, majority of the traditional (e.g. Biguanides, Insulin, Statins, RAAS inhibitors, β-blockers, diuretics) and emerging (e.g. SGLT2 inhibitors, GLP-1 receptor agonists, DPP-IV inhibitors, P2Y12 antagonists) medications for managing DCM have several adverse effects including, hypotension, urinary tract infections, diabetic ketoacidosis, pancreatitis, gastrointestinal malaise, renal impairments, and hepatotoxicity^9^. <u>Therefore, there is a need for safer, effective and novel therapeutic modalities with promising validated molecular targets related to pathophysiological mechanisms contributing to DCM.</u>

EphB tyrosine kinase family members have different roles in cardiovascular-related diseases^10^. For example, genes encoding Eph receptor tyrosine kinases, particularly EPHB1, were significantly downregulated in cardiomyocytes of hypertrophic hearts, and silencing EPHB1 expression resulted in hypertrophy of cardiomyocyte^11^. EphrinB1 knockout mice show diastolic dysfunction that progresses to systolic heart failure and 100% mortality at adulthood^12,13^, and heart failure with preserved ejection fraction (HFpEF) patients have downregulation of Efnb1 in ventricles^12^, suggesting that drugs that selectively inhibit EphB1 in the heart may have limited therapeutic potential. In contrast, EphB2 expression in upregulated in mouse models of cardiac hypertrophy and in patients with cardiac fibrosis^14^, whereas, deletion of EphB4 in endothelial cells lead to cardiac capillary rupture in mouse models^15^. Despite such evidence, there are no approved drugs that specifically target EphB signaling to treat DCM. Recently, we reported pan-EphB tyrosine kinase inhibitors sparing EphB3 in metabolic syndrome termed STA-analogs, where STA-013 induced whole-body weight loss while preserving the lean muscle mass, restored glucose homeostasis, and mitigated insulin resistance in diet-induced obese (DIO) mouse models^16^. Our objectives were to test the efficacy of STA-013 on glycemic control, weight loss, and indices of cardiac function and fibrosis ina preclinical mouse model of high-fat diet (HFD) and streptozotocin (STZ)-induced diabetic cardiomyopathy.

## Results and Discussion

We assessed STA013-mediated effect on diabetic cardiac dysfunction in STZ-HFD mouse model of diabetic cardiomyopathy (**Figure 1A)**. STA-013 treatment (25 mg/kg BW) resulted in a significant reduction in whole-body weight (**Figure 1B**), accompanied by improved glucose homeostasis (**Figure 1C and 1D**), and enhanced insulin sensitivity (**Figure 1E and 1F**). Interestingly, echocardiographic analysis revealed marked improvements in cardiac function, including ejection fraction (**Figure 1G and 1H**) and fractional shortening (**Figure 1I**) with no changes in heart weight to body weight (HW/BW) ratios, compared to vehicle-treated STZ-HFD mice. The cardiac improvements by STA-013 are unlikely to be EphB1 mediated because EphrinB1 knockout mice have diastolic dysfunction^12,13^, and EphrinB1 in downregulated in HFpEF patients^12^. Similarly, EphB4 inhibition by STA-013 is unlikely to exert a major effect as specific deletion of EphB4 in endothelial cells lead to cardiac capillary rupture in mouse models^15^. However, the cardiac improvements by STA-013 are very likely to be mediated by EphB2 because others have shown that EpHB2 is overexpressed in preclinical mouse models of myocardial infarction and cardiac hypertrophy, as well as in patients with severe cardiac fibrosis with a concurrent increase in fibroblast proliferation, myofibroblast activation, and collagen deposition, and EphB2 knockdown improved ejection fraction in myocardial infarcted mice^14^. Next, we harvested the hearts for Masson’s trichrome staining and found a significant reduction in myocardial collagen deposition in STA013-treated mice compared to vehicle controls, indicating attenuation of cardiac fibrosis (**Figure 1S and 1T**). Additionally, immunoblotting revealed that STA-013 inhibited phosphorylation of EphB tyrosine kinase forward signaling in the hearts isolated from the mice-treated with STZ and HFD, compared to vehicle-treated STZ+HFD mice (**Figure 1U and 1V**). This was associated with significant increase in insulin receptor beta (IRβ) signaling, and a decrease in TGF-β levels, in STA-013 treated-STZ-HFD mice, compared to vehicle treated-STZ-HFD mice (**Figure 1W, 1X, 1Y**). These findings further support the notion that the effects of STA-013 are potentially mediated *via* EphB2 signaling because EphB2 contributed to cardiac fibrosis through activation of the Stat3 and TGF-β/Smad3 signaling, and EphB2 knockdown decreased the fibrotic scar size in mice with myocardial infarction, compared to controls^14^. However, the specificity of STA-13 action needs to be confirmed with EphB specific gain and loss of function models.

**Figure 1:**
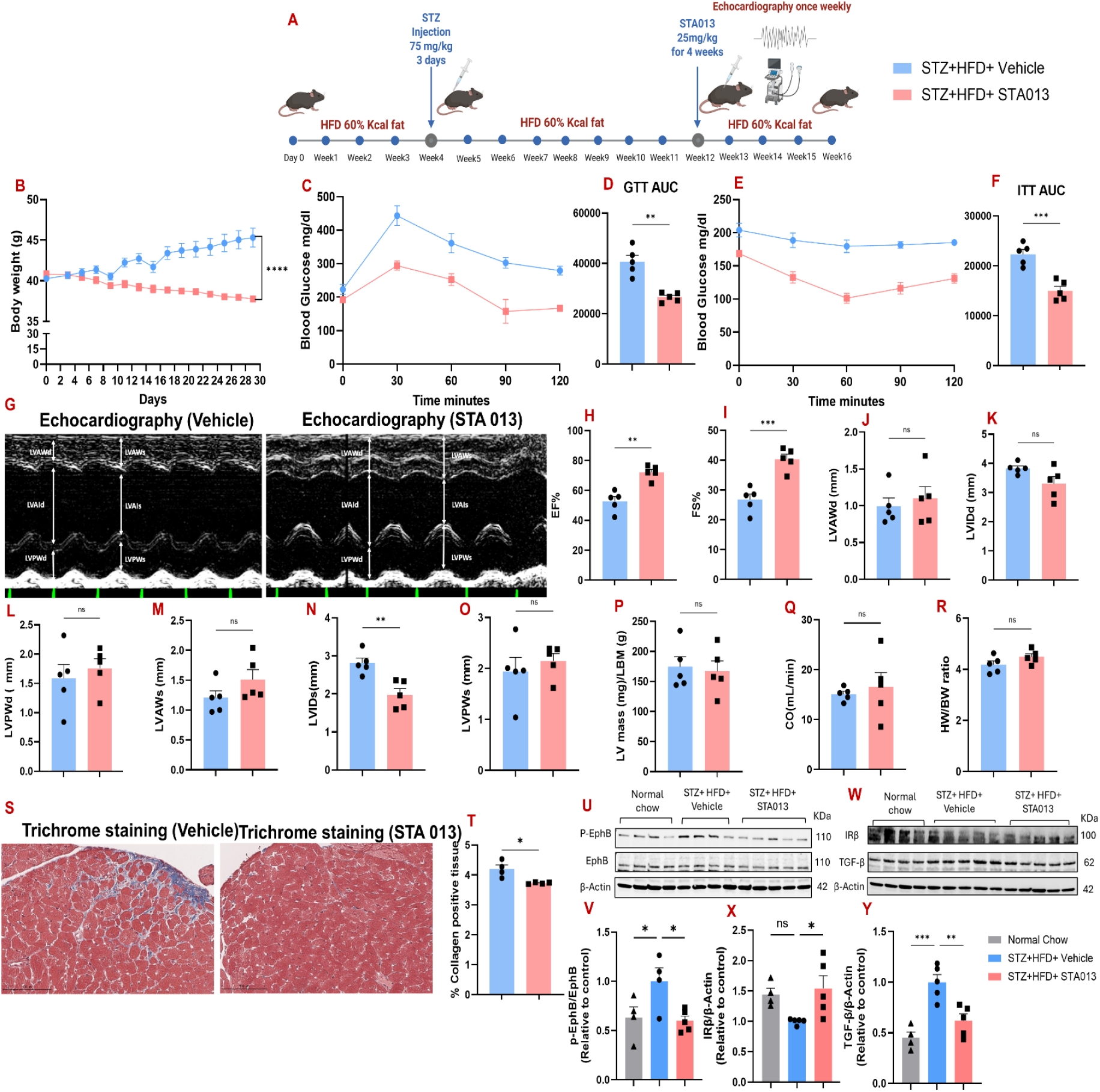
Pharmacological Inhibition of EphB Signaling by STA-013 Improves Metabolic Homeostasis, Cardiac Function, and Reduces Fibrosis in STZ-HFD–Induced Diabetic Cardiomyopathy. **A.** Schematic for *in vivo* administration of STA 013 in STZ-HFD-induced DCM mouse model. **B**. Body weight measurement over 30 days of administration of vehicle and STA 013 (25mg/kg/day, IP) (n=5 mice). **C and D**. Glucose tolerance test and AUC data. **E and F**. Insulin tolerance test and AUC data. **G**. Representative echocardiography M-mode images for vehicle and STA 013 treated mice with DCM. **H**. Ejection fraction percentage (EF%). **I**. Fractional shortening percentage (FS%). **J**. Left ventricle anterior wall in diastole (LVAWd). **K**. Left ventricle internal diameter in diastole (LVIDd). **L**. Left ventricle posterior wall in diastole (LVPWd). **M**. Left ventricle anterior wall in systole (LVAWs). **N**. Left ventricle internal diameter in systole (LVIDs). **O**. Left ventricle posterior wall in systole (LVPWs). **P**. Left ventricle (LV) mass/lean body mass (LBM). **Q**. Cardiac output (CO). **R**. Heart weight (HW) to body weight (BW) ratio. **S**. Representative images of Masson’s trichrome staining of heart sections showing fibrotic tissue for vehicle and indacaterol treated mice with DCM. **T**. Collagen deposition positive tissue (%). Data is presented as mean ± SEM, statistical significance was analyzed using a student’s t-test; where ****p<0.0001, ***p<0.001, **p<0.01, and ns nonsignificant. **U and W**. Immunoblots for heart tissue for vehicle and STA 013 treated mice with DCM. Densitometry analysis for **V**. p-EphB/EphB, **X**. insulin receptor beta (IRβ), and **Y**. transforming growth factor beta (TGF-β). Data is presented as mean ± SEM and analyzed by one-way ANOVA, followed by Bonferroni post-post hoc test; where ***p<0.001, **p<0.01, *p<0.05 and ns non-significant.

## Conclusions

In summary, we previously reported STA-013 as a pan-EphB tyrosine kinase inhibitor that restored glucose homeostasis and mitigating insulin resistance. We expand on these findings and now show that STA-013 protected against cardiac dysfunction and fibrosis in HFD-STZ mice model. Taken together, these results suggest that STA-013 not only improved systemic metabolic functions associated with T2D but also directly mitigated structural and functional deterioration of the diabetic heart.

## Materials and Methods

### Animals

Ethical approval for all animal-related activities was obtained from the Institutional Animal Care and Use Committee at Texas Tech University Health Sciences Center. All mice were kept under pathogen-free conditions, 12-h light-dark cycle, controlled temperature (20–22°C), and fed either a normal chow diet (70 % CHO, 20 % proteins, and 10 % fats by kcal) or high-fat diet (60 % fats by kcal Research Diets Inc, D12492) and water ad libitum. Mice were obtained from Jackson lab (C57BL/6 J 000664). 8-week-old male C57BL/6J mice were fed HFD (60% fat kcal, Research Diets®) for 12 weeks to develop obesity, and in week 4 will be administered with a low-dose of streptozotocin (75 mg/kg, IP for 3 consecutive days) to induce DCM.

### Glucose and insulin tolerance test

Glucose and insulin tolerance tests were performed in mice fasted overnight for glucose tolerance test and after a 4 h fast for insulin tolerance test, following which IP glucose (1 g/kg) or insulin (0.5 U/kg) was administered. Blood glucose measurements were assessed via tail whole-blood at the end of the fast (0 min), followed by samples at 15, 30-, 60-, 90-, and 120-min post-glucose or insulin administration using a blood glucose monitoring system, as previously reported^16^.

### Echocardiography of Mice

Assessment of in vivo heart function on sedated mice were performed following our published procedures^17–19^. Briefly under isoflurane anesthesia, echocardiography was performed using a 10-MHz probe (Prospect T1, Scintica Inc®) at baseline, and at 1, 3, 5, and 7 weeks after start of drug administration. Echocardiographic M-mode images were obtained from a parasternal short-axis view at the level of the papillary muscles. Left ventricular internal diameters at end-diastole (LVIDd) and end-systole (LVIDs) will be measured from M-mode recordings. Six representative contraction cycles were selected for analysis, and average indexes (LVIDd, LVIDs, left ventricular ejection fraction (LVEF), fractional shortening, E/A) were calculated for each mouse. All echocardiography measurements were performed in a blinded manner.

### Immunoblotting

Immunoblotting of heart tissue were performed following our published procedures^17–19^. Briefly, after extraction of protein from heart tissues, total protein concentration was quantified, proteins were transferred to PVDF or nitrocellulose membranes, blocked in 5 % BSA/TBST, and membranes incubated with commercially available (Thermofsiher, Cell Signaling, Santa Cruz, Abcam) primary antibodies for targets in Ephrin signalling (anti-EphB2, anti-phospho-EphB Pan), insulin signaling (anti-INSR β), growth factor signaling (TGF-β, P-Smad3 (Ser423/425), Smad3), and with internal loading controls (anti-β-actin). Horseradish peroxidase-conjugated anti-rabbit (Cell signaling 7074, 1:2000) or Alexa Flour 680 goat anti-rabbit (Thermofisher Scientific A21109, 1:5000) will be used as secondary antibody and incubated for one hour at room temperature. The membranes were incubated with SuperSignal West Femto (Thermofisher) chemiluminescent substrate for 5 mins and then explored using LICOR Odyssey Fc for chemiluminescent or fluorescence detection and quantified by image studio software version 5.5.

## Acknowledgments

This research was funded by start-up funds offered by the School of Pharmacy at TTUHSC and the Office of Innovation and Research at TTUHSC, awarded to Mahmoud Salama Ahmed. Funding support from the American Heart Association (Grant # <u>953881</u>), Diabetes Action (Grant# <u>523</u>) and Texas Tech University School of Veterinary Medicine to Prasanth K. Chelikani is acknowledged.

